# Optimization of cryopreservation and *in vitro* fertilization techniques for the African turquoise killifish *Nothobranchius furzeri*

**DOI:** 10.1101/2020.04.14.040824

**Authors:** Luca Dolfi, Tsz Kin Suen, Roberto Ripa, Adam Antebi

**Affiliations:** Max Planck Institute for Biology of Ageing, Cologne, Germany; Cologne Excellence Cluster on Cellular Stress Responses in Aging Associated Diseases (CECAD), University of Cologne, Cologne, Germany

## Abstract

Over the last decade, the African turquoise killifish, *Nothobranchius furzeri*, has emerged as an important model system for the study of vertebrate biology and ageing. However, rearing this fish in captivity can pose challenges, due to the short window of fertility, inbreeding problems, and the continuous maintenance of different strains and transgenic lines. To date, the main means of long term strain maintenance is to arrest embryos in diapause, a poorly understood and unreliable method. To solve these problems, we developed a robust protocol to cryopreserve sperm and to revive them for *in vitro* fertilization (IVF), as a better option for long term storage of *N. furzeri* lines. We tested a variety of extender and activator buffers for sperm *in vitro* fertilization, as well as cryoprotectants to achieve maximal long term storage and fertilization conditions tailored to this species. Our optimized protocol was able to preserve sperm in a cryogenic condition for months and to revive an average of 40% upon thawing. Thawed sperm were able to fertilize nearly the same number of eggs as natural fertilization, with an average of ~25% and peaks of ~55% fertilization. This technical advance will greatly facilitate the use of *N. furzeri* as a model organism.

## Introduction

Over the last few years the African killifish, *Nothobranchius furzeri,* has emerged as important model system for the study of vertebrate ageing. The life cycle of these fish is characterized by a fast growth rate, reaching sexual maturity by 4-5 weeks, and a maximum lifespan of 6,5-7 months (as mean lifespan of the most long-lived 10% of a given cohort)[1], [2], making them among the shortest-lived vertebrate species bred in captivity and a unique platform for the rapid exploration of aging and age-associated diseases[3].

Unfortunately fast growth and aging features also carry some drawbacks that make the maintenance in captivity of this species quite challenging. Rapid growth and maturation lead to a rapid passing of successive generations, which can often result in inbreeding [4], [5]. Indeed, it is virtually impossible to establish an ancient founder to maintain an original genotype because of the marginal overlap of generations. Fertility in this species is limited to the 5^th^-20^th^ week of life and is maximal between the 6^th^ and the 11^th^ [6]–[8], marking an extremely narrow breeding window, especially compared to other fish species where this can correspond to years. Moreover the success of transposon-mediated transgenesis [9] and CRISPR-mediated mutagenesis [10] in this fish means that more genetically engineered lines require continuous maintenance and space usage. Breeding to preserve a line takes considerable effort and is fraught with risk of accident or infection that can result in strain loss. Furthermore, *N. furzeri* husbandry requires a large amount of space, since this species is optimally grown in captivity when single housed because of fish-to-fish harassment and food competition [8].

Given these constraints, it is essential to develop protocols to maintain stocks without constant breeding. Currently, the only known way to “freeze” a generation is through diapause, a state of arrested development [2]. However, diapause itself is quite variable and a topic of intense investigation [11], [12]. It is difficult, to date, to induce and release diapause in a controlled and synchronized manner from a large pool of embryos. As well, it is challenging to retain permanence in this stage. Even eggs in diapause need periodic maintenance, their medium or substrate must be checked, cleaned and changed. As the number of strains, species or lines to preserve increases, this method becomes quickly untenable at larger scales.

To solve these problems, researchers of other fish species rely on *in vitro* fertilization and sperm cryopreservation techniques. Through cryopreservation, it is possible to create sperm banks that can store fish genetic pools with minimum maintenance effort for years. Upon revival, the thawed sperm can usually fertilize eggs with a success rate from10-80% [13]. Specific protocols have been established to preserve and activate the sperm of both production fish species (salmonids, sturgeons, carps and catfishes) and research species *(Zebrafish, Medaka)* [14]. Despite this, there is no protocol available to date for *in vitro* fertilization or sperm cryopreservation of any killifish species.

Here we optimized a protocol for Killifish sperm cryopreservation and *in vitro* fertilization. Our protocol will greatly facilitate the husbandry and the usefulness of *N. furzeri* as a model organism for research.

## Results

### Extender and Activator

The optimum conditions for a suitable fertilization environment can vary greatly for each fish species, as well as the protocols, buffers and setups for spermatozoa cryopreservation. For most freshwater fish, sperm motility can be initiated by hypotonic osmolalities [15] and/or by alteration of ion concentration such as potassium or calcium [16], [17]. Once activated, the sperm usually have a short period of motility (30 s to 5 min), depending on species [15].

Protocols from other fish species avail of an “extender” and an “activator” solution. The extender is usually a saline-buffered solution that is mixed with the extracted sperm that keeps it in a stable and inactive condition. This is made possible mostly by the high molality of the extender solution, which is comparable or higher than the ion concentrations inside the gonad [13], [18].

By contrast, the activator is a low molality solution, which when added to the sperm-extender solution, initiates sperm activity and movement. Often but not always the activator is a dilution of the extender and specific concentrations of ions like potassium, calcium and magnesium are required for proper activation [16]. Thus, aside from osmolality, medium composition is also responsible for correct sperm motility, directionality and endurance [13], [19].

For the most common laboratory species like *Zebrafish* and *Medaka* there are several existing protocols that describe the use of Hank’s Balanced Salt Solution (HBSS), Buffered sperm motility-inhibiting solution (BSMIS) or Fetal bovine serum (FBS) as extender and a dilution of these compounds or the addition of another composite salt, like Instant Ocean or Iwamatsu solution, for activation [14], [20], [21].

Our first goal therefore was to test sperm activation upon mixing with different known solutions. The sperm was collected from male fish through dissection and gonad isolation. Before dissection, the fish were carefully dried to prevent spontaneous sperm activation due to water presence. Both testes were placed into liquid solution in an Eppendorf tube and quickly shaken for 5-10 seconds to allow the sperm to be released in the medium. We tested different activating solutions including Tank water, Deionized water, BSMIS, FBS, HBSS and Iwamatsu solution at different dilutions and molalities (Fig 1).

**Fig 1:**
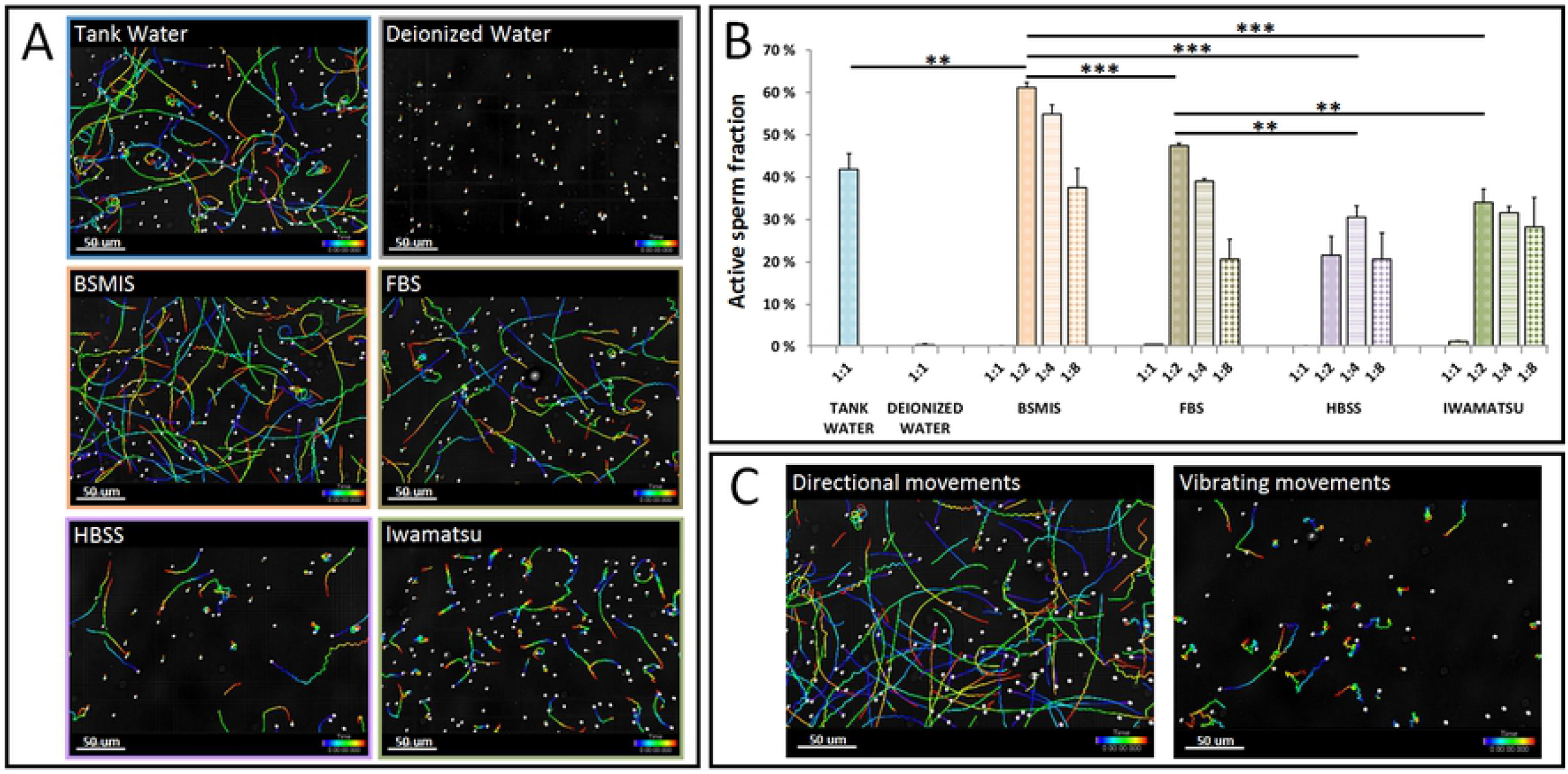
Fresh sperm activation in different buffers. (A) Tracking imaging of sperm movements. (B) Quantification of sperm activation in different buffers. (C) Magnification of different types of movement occurring after sperm activation, proper directional movements on the left and erratic inefficient vibrating movements on the right. Asterisks indicate significance levels of between-condition t-test comparisons. *p < 0.05, **p < 0.01, ***p < 0.001. N = 3 biological replicates for each condition.

Immediately afterwards, the gonads were removed from the medium, 10 μl of the mixture was put in a hemocytometer chamber and sperm movements were video-recorded under a microscope. The videos were subsequently analyzed and the percentage of directionally moving spermatozoa was estimated (Figs 1A and 1B).

Tank water, the natural medium where fish biologically breed, was able to activate ~40% of the sperm. Deionized water was able to trigger a very initial activation for a few seconds (data not shown), which faded immediately thereafter, resulting in the appearance of either vibrating or immobile sperm (Fig 1C). BSMIS, FBS, HBSS and Iwamatsu solutions were able to activate sperm only when diluted, while the stocks solutions molalities were too high to promote sperm activation (Fig 1B). Notably BSMIS and FBS diluted to 1:2 and 1:4 had comparable or even superior activation rates compared to tank water while HBSS and Iwamatsu solution instead were inferior as activation promoters at any dilution.

Aside from dilution of the extender itself, dilution with FBS is often used in combination with other solutions for sperm activation [20], [22], [23]. For example, Aoki et al. used FBS combined with 2 more volumes of Iwamatsu solution to promote directional activation in *Medaka* (which is the closest species to *Nothobranchius* among model fish). We therefore combined the FBS buffer with other solutions as candidate activators in order to achieve an increased yield of activation. We tested FBS in combination with two volumes of Iwamatsu, BSMIS and HBSS, at different dilutions (Fig 2A).

**Fig 2:**
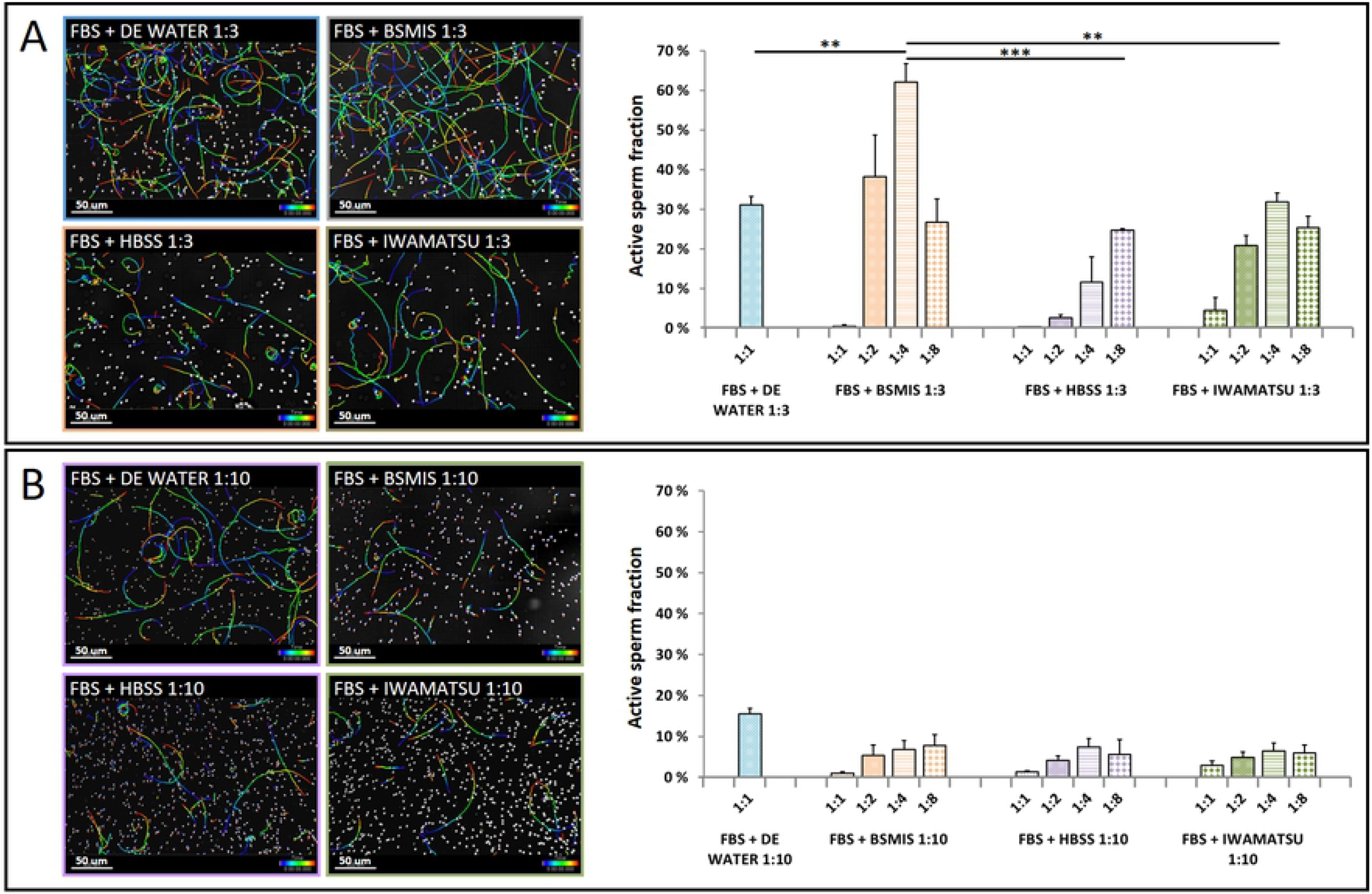
Frozen sperm activation in different buffers at different ratios. (A) Tracking imaging of sperm movements and quantification of directional sperm in different activating solutions, mixed in a ratio 1 to 3 with the extender solution (1 volume of extender solution in 3 volumes of activating solution). (B) Tracking imaging of sperm movements and quantification of directional sperm in different activating solutions, mixed in a ratio 1 to 10 with the extender solution (1 volume of extender solution in 10 volumes of activating solution). Asterisks indicate significance levels of between-condition t-test comparisons. *p < 0.05, **p < 0.01, ***p < 0.001. N = 3 biological replicates for each condition.

In contrast to *Medaka,* FBS mixed with Iwamatsu did not give remarkable activation yields, probably because of the high molality of the resulting mixture, which may be suitable for *Medaka* (a half saltwater half freshwater fish) but not for freshwater *N. furzeri.*

Among all the combinations, we noticed an impressive activation when applying FBS together with a dilution 1:4 of BSMIS, reaching activation yields >60%. These results outperformed all the other combinations and the baseline activation in tank water, being similar or superior to the yields achieved by BSMIS 1:2 and BSMIS 1:4 obtained in the previous test (Figs 2A and 1B). As a second test, we repeated the previous experiment changing the ratio between the extender (FBS) and the activators, from 1 plus 2 volumes respectively to 1 plus 9 volumes respectively. However, the activation rates obtained in the second test (Fig 2B) were significantly worse compared to the previous test and in these conditions most of the spermatozoa were vibrating or not moving at all.

### Cryoprotection, Freezing and Thawing

To allow the sperm to survive the freezing procedure and be preserved for a long time in a cryostatic condition, the extender has to be supplemented with a cryoprotectant. Among the most used are DMSO, DMF, MetOH glycerol and DMA, with a concentration that varies between 5 and 20% (usually 10%) of the volume [14], [24]. These chemicals, even if mildly toxic, have the property to surround the sperm cells, preventing the formation of ice crystals that could compromise membrane integrity and protecting them from cryodamage [25].

Thus, our second goal was to establish conditions for long-term sperm cryopreservation. Our target was to find an extender solution, which combined with the proper cryoprotectant, could preserve the collected sperm in the inactive state in a frozen condition. Since BSMIS and FBS worked best to activate sperm when diluted (Fig 1B), we focused our studies on these.

In the first experiment we mixed the selected extender with an arbitrary concentration (10%) of DMSO, DMF, MetOH, DMA or glycerol for a total of 10 combinations. Sperm was released in already mixed extender-cryoprotectant solutions (Fig 3C) and the mixture was incubated for 1h at 4°C (Fig 3D), allowing the cryoprotectants to be evenly absorbed by the sperm cells.

**Fig 3:**
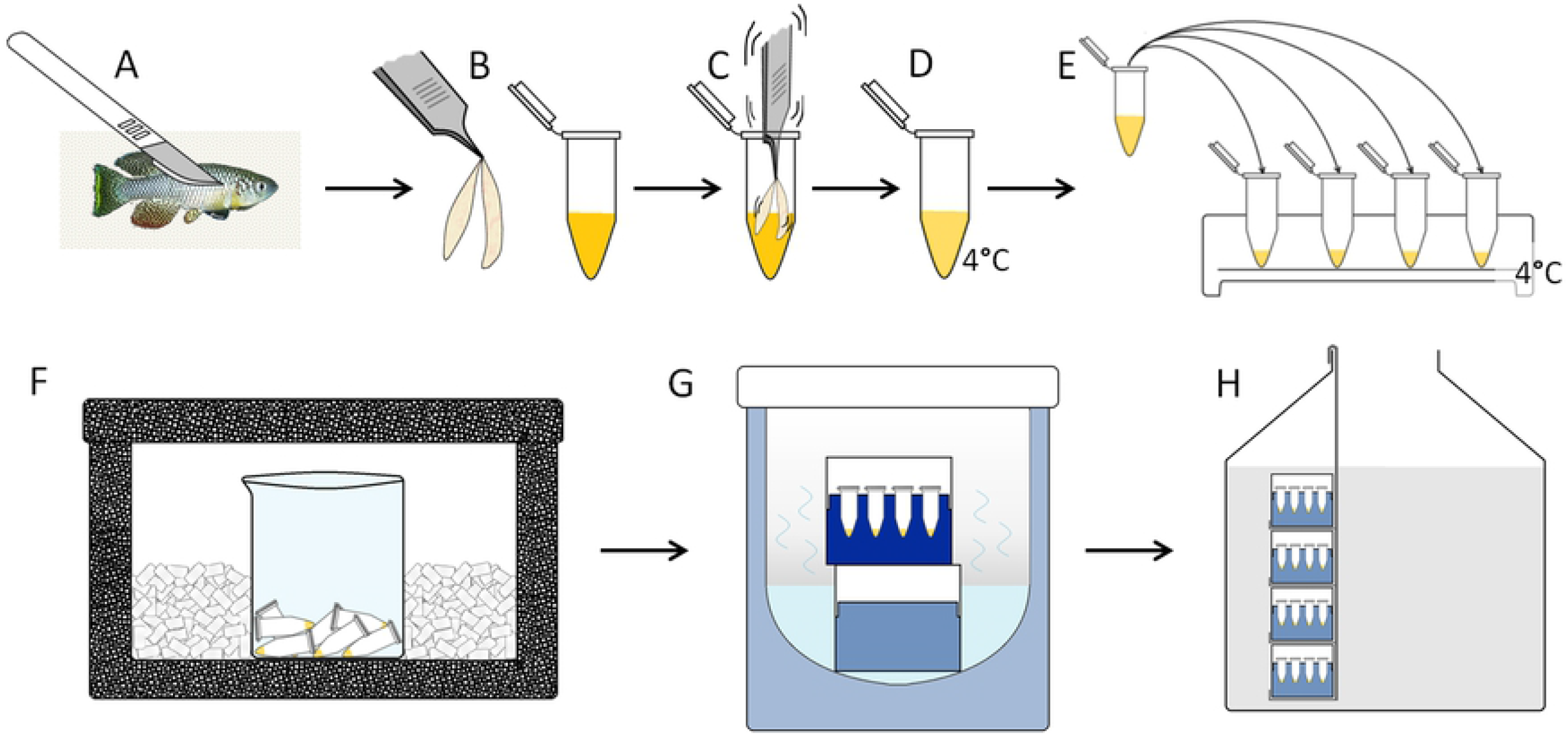
Schematic of sperm collection and freezing procedure. (A) Males dissection. (B) Gonads extraction. (C) Gonads swing in extender plus cryoprotectant solution. (D-E) Cryoprotectant effect on sperm cells at 4°C and solution transfer in smaller aliquots. (F) First freezing step setup with laying on the bottom of a glass beaker surrounded by dry ice in a closed styrofoam box. (G) Second freezing step setup with aliquots exposed to nitrogen vapor. (H) Aliquots long term storage in liquid nitrogen. For a detailed explanation of the several steps refer to the text.

The samples were then aliquoted in 60ul volumes (Fig 3E), frozen in dry ice (Fig 3F), and nitrogen gas phase (Fig 3G) in sequence, and finally stored in liquid nitrogen (Fig 3H).

After 24–48h freezing, the sperm were revived through fast thawing in a 30°C water bath and activated by the addition of 2 volumes of BSMIS diluted 1:4 in the case of FBS extender and BSMIS diluted 1:2 in the case of BSMIS extender. We found that sperm cryopreserved using 10% DMSO as cryoprotectant maintained the highest activation (Fig 4A), followed by 10% methanol (with a greater effect when used in combination with FBS), and last with DMF.

**Fig 4:**
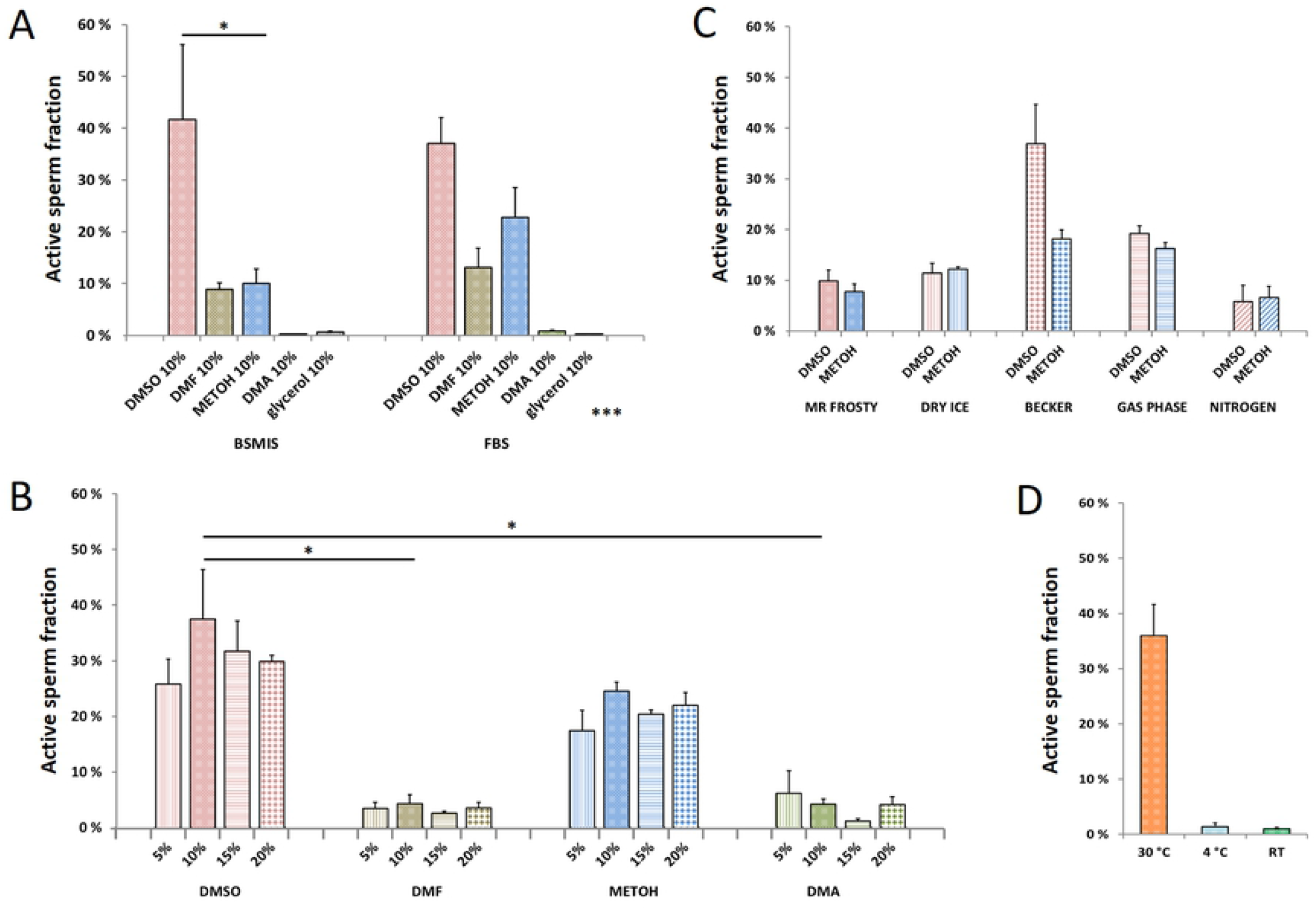
Cryoprotectants, cryopreservation and thawing efficiencies. (A) Frost protecting efficiencies of different cryoprotectants in different extender solutions. Y bar represents the portion of sperm able to reactivate upon defrosting and mix with the activator solution. (B) Different concentrations effects of cryoprotectants in FBS extender. (C) Different freezing methods with different cooling rates applied to FBS extender plus 10% cryoprotectant. (D) Different thawing methods with different thawing rates applied to FBS extender plus 10% DMSO. Asterisks indicate significance levels of between-condition t-test comparisons. *p < 0.05, **p < 0.01, ***p < 0.001. N = 3 biological replicates for each condition.

Glycerol and DMA were absolutely ineffective in protecting the sperm from freezing, resulting in immobile or inviable sperm upon thawing.

To achieve a higher yield of sperm survival and activation, we further optimized the cryoprotectant concentration in the sperm-extender mixtures. Several concentrations of cryoprotectants were tested in combination with FBS. Sperm cryopreserved in FBS with 10 to 20% DMSO maintained comparably high activation (Fig 4B), dropping slightly on higher concentrations, probably due to increasing toxicity. Any concentration of methanol was able to moderately protect sperm from cryodamage even though the revival rates were inferior compared to DMSO by 5% to 20% less. Other combinations were not able to protect sperm cells efficiently. Less than 10% of sperm were able to revive after freezing in any concentration of DMA or DMF (Fig 4B).

Since no relevant increase in sperm survival and activation was observed in the samples with a cryoprotectant concentration >10%, we set this concentration as a standard for our future cryopreservation experiments. We selected DMSO and Methanol as cryoprotectants to perform further optimization of cryopreservation.

Apart from the application of cryoprotectants, the freezing procedure is also a crucial step for cryopreservation. A slow freezing rate can produce large ice crystals and damage cellular ultrastructure, whereas a rapid freezing rate induce only small intracellular ice crystals that are less likely to prompt damage [26].

We optimized sample freezing rate with various freezing setups. We placed the Eppendorf tubes in 1) Mr. Frosty™ Freezing Container (estimated −10°C per min), 2) a glass beaker surrounded by dry ice (estimated −20°C per min), 3) direct contact with dry ice (estimated −50°C per min), 5) a Dewar vessel calorimeter partially filled with liquid nitrogen, where the Eppendorf tubes were placed in a box without direct contact with liquid nitrogen but exposed to nitrogen gas phase (estimated −100 °C per min); and 6) direct liquid nitrogen contact (estimated −200°C per min) (Fig 4C). Sperm vials from all freezing setups were stored in liquid nitrogen overnight after they reached a temperature below −50°C. After one day, frozen sperm samples were revived using BSMIS 1:4 and observed for their activation. Sperm vials frozen in a beaker surrounded by dry ice (±20°C per min) achieved the highest activation upon revival. Sperm cryoprotected with 10% DMSO had higher activation than that cryoprotected with 10% methanol (Fig 4C), similar to above. Therefore, we selected DMSO as the final cryoprotectant for our cryopreservation protocol.

To assess the proper thawing rate for the frozen sperm vials, we thawed the vials at 1) 30°C in a water bath, 2) room temperature and 3) 4°C in the fridge to achieve rapid, medium, or slow thawing rates, respectively. We then revived sperm using BSMIS 1:4 and observed the activation rate. Our results showed that rapid thawing achieves the highest survival and activation of sperm (Fig 4D).

### Egg fertilization and survival

Finally, we tested if the active sperm obtained with our protocol was also able to fertilize eggs obtained from *N. furzeri* females. To perform this experiment in the most comparable way, we set up several aquarium tanks with 5 female and 2 male specimens per tank and allowed them to naturally breed for 2 days. We then collected the eggs generated from the natural breeding and monitored their survival rate until mid somitogenesis. After the natural breeding, the males were separated from the females, kept alone for 2 days, then sacrificed for gonad extraction. The females were anesthetized, dried carefully (Fig 5A), and their unfertilized eggs were gently pushed out from their belly (Fig 5B).

**Fig 5:**
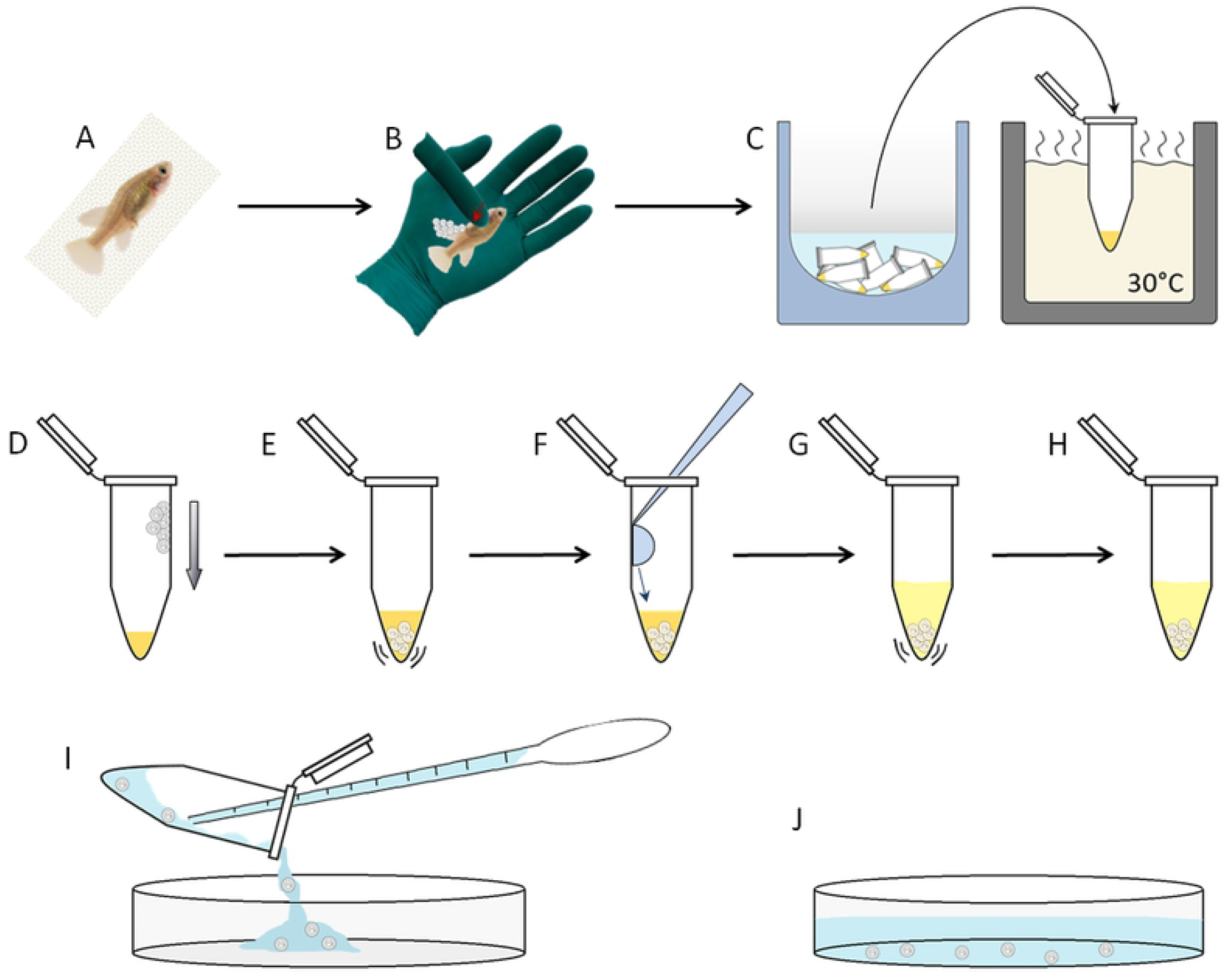
Schematic of sperm thawing and egg fertilization procedure. **(A)** Females preparation. (B) Eggs collection through gentle belly squeeze. (C) Frozen sperm thawing in a water bath. (D-E) Eggs mixing with sperm-extender solution. (F) Activating solution addition and (G) mix. (H) Fertilization occurring. (I) Eggs recovery in a petri dish. (J) Fertilized egg developing in a petri dish. For a detailed explanation of the several steps refer to the text.

The sperm obtained from the male gonads was mixed with FBS or BSMIS buffer with 10% DMSO. Half of the sperm-extender-cryoprotectant mixture was directly activated (with 1:2 or 1:4 BSMIS) and used for the fertilization of the eggs (Fig 5D-J) while the other half was frozen and cryopreserved (Fig 3).

After a variable time period (from 1 day to 2 months), the frozen sperm were thawed, activated and used to fertilize another pool of eggs, following the same procedure (Fig 5).

All the eggs fertilized in both ways were monitored until the stage of mid somitogenesis (Fig 6C) or later (Fig 6D) and some up to the point of hatching (Fig 6E). Survival rates were scored (Fig 6F).

**Fig 6:**
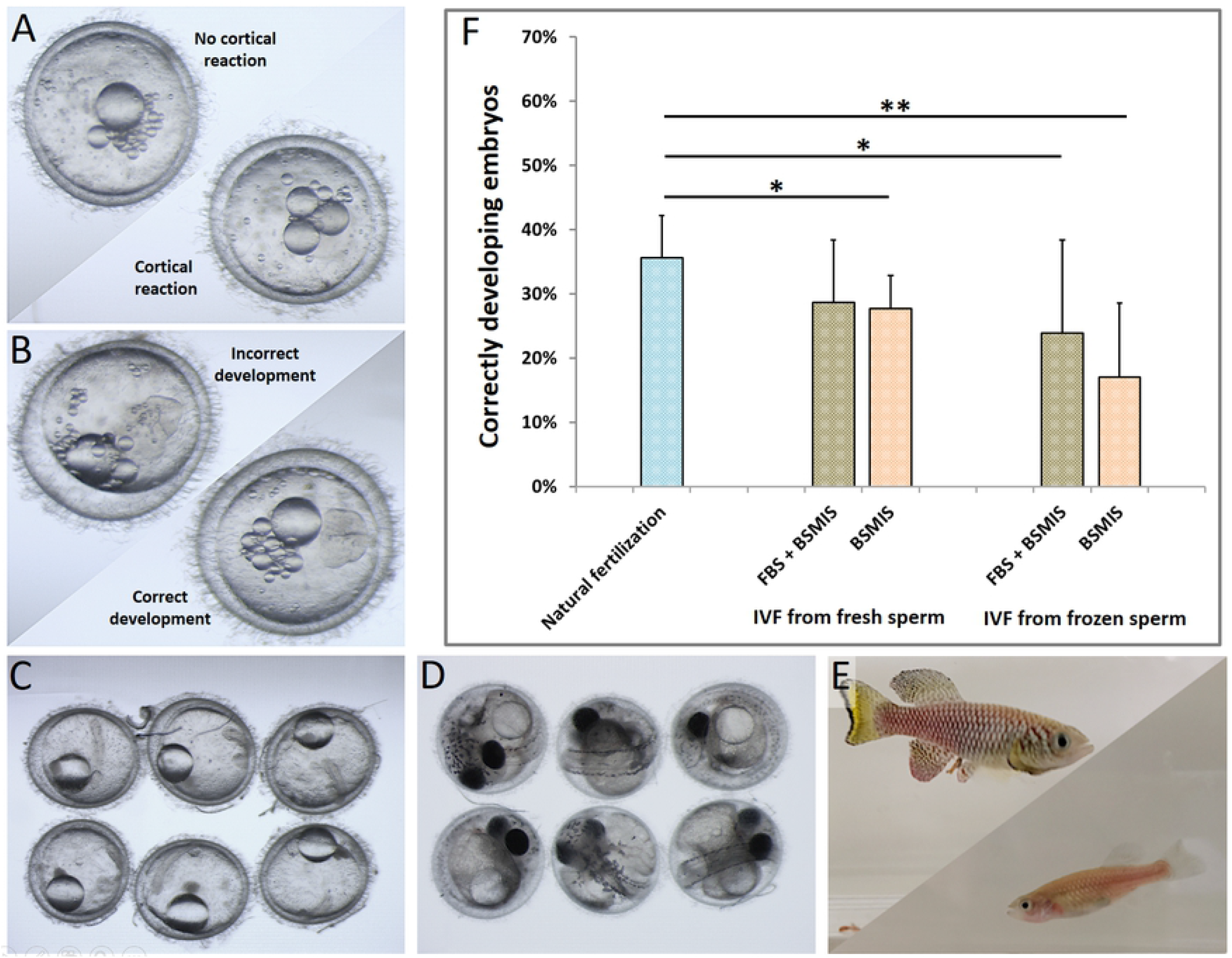
Fertilization rates and embryos development. (A) Embryos with not occurred (top left) and occurred (bottom right) cortical reaction. (B) Embryos developing not correctly with odd cells number with different size (top left) and correctly with a proper 4 cells stage (bottom right). (C) Embryos fertilized with IVF developing through diapause II / mid-somitogenesis stage and later at (D) and advanced developmental stage. (E) Adult fish derived from IVF embryos. (F) Fertilization efficiencies of natural breeding, IVF with fresh extracted sperm and IVF with frozen sperm. Asterisks indicate significance levels of between-condition t-test comparisons. *p < 0.05, **p < 0.01, ***p < 0.001. N = 6,8,4,10,3 biological replicates for each fertilization condition, respectively, in order of appearance in the graph.

For the detailed fertilization protocol, females were anesthetized and carefully dried from any residual water to prevent spontaneous egg cortical reaction or activation (Fig 5A). Laying the fish on an open hand, a gentle pressure was applied with a finger on the female belly, pushing gently from the middle toward the anus (Fig 5B). 5 to 35 eggs were usually expelled. Those eggs were collected using forceps in an Eppendorf tube containing the extender-sperm solution (Fig 5D). In the case of frozen sperm, the tube was thawed immediately before in a 30°C water bath (Fig 5C). These actions were performed by two people, with one person thawing the sperm as soon as the other one began to expel eggs from the female. Eggs were placed at the edge of the tube and gently pushed to the bottom (Fig 5D). The best yields of fertilization were achieved using aliquots of sperm-extender of 60 ul and no more than 35 eggs per aliquot. Once the eggs were completely immersed, the tube was gently flicked for 10-20 seconds (Fig 5E), allowing the mix to homogeneously distribute around all the eggs. The activator solution was then pipetted into the tube letting drops slide over the tube’s border (Fig 5F) and then mixed with the extender by gently flicking the tube for 20-30 seconds (Fig 5G). The activated sperm was left with the eggs for 10 minutes and the tubes standing open on a bench at room temperature (Fig 5H). At this step, 10ul of the mixture were pipetted under the microscope to evaluate sperm motility. To avoid damage due to DMSO toxicity, the embryos were transferred after 10 minutes to a petri dish using a pipette and methylene blue buffered tank water (Fig 5I). The water was replaced twice and the petri dish incubated at 28°C (Fig 5J).

We noticed that the cortical reaction, the earliest process of development where the distance between the yolk and chorion membrane increases, was not correlated with fertilization success in *N. furzeri.* Once in contact with an aqueous medium, a large number of eggs were able to spontaneously undergo the cortical reaction in the absence of sperm, while a smaller percentage remained blocked in the pre-cortical reaction stage (Fig 6A).

This happened also during natural fertilization, where a small fraction of collected eggs was found blocked in the pre-cortical reaction stage. Eggs that underwent a spontaneous cortical reaction showed abnormal cell cleavages, errant development and embryonic death in the first 10 days (Fig 6B). Analogously, eggs unable to carry out the cortical reaction inevitably led to an unsuccessful fertilization, even in the presence of sperm and the proper buffer. Though the mechanism underlying the cortical reaction remains unknown in this species, it is most likely related to the degree of egg maturation [27], [28]. Our analyses were thus made by counting the total number of embryos developed until mid somitogenesis (Fig 6C) over the total initial number of eggs used for fertilization (Fig 6F). Moreover, several embryos were allowed to develop and were followed post-somitogenesis (Fig 6D), post-hatching and up to adulthood (Fig 6E). The growth rates were normal and no defects were detected. Those fish were fertile and able to produce viable embryos.

In conclusion, we found that under our conditions, rates of fertilization with frozen sperm ranged from 15 to 25% and were only slightly below fresh IVF or natural fertilization. FBS and BSMIS were better at *in vitro* fertilization than BSMIS alone (Fig 6F). A detailed protocol for the entire procedure is found in the Supplemental methods.

## Conclusion and Discussion

In this study we optimized sperm activation, cryopreservation and *in vitro* fertilization in the species *N. furzeri,* aiming to establish protocols to obviate problems linked to inbreeding, line maintenance and space usage. Previous publications show how an osmolality drop together with a species specific salt combination are required for fish sperm activation. Concentrating on the most used buffers and testing them in several dilutions and combination, we found that *N. furzeri* requires a specific protocol tailored for this species, which represents a hybrid between *Medaka* and *Zebrafish.*

After exploring several combinations, we narrowed down the viable extender solutions to FBS and BSMIS and activators to BSMIS 1:4 and 1:2. Concerning cryoprotectants only DMSO and MetOH achieved remarkable results in protecting against cryodamage, DMF in very few cases, while DMA and glycerol not at all.

Besides the choice of the extender, activator and cryoprotectant, there were other critical variables that drastically influenced the outcome of our IVF. The proper growth, health, and age of the fish were among the most important features influencing sperm and egg quality. Sperm or egg pools derived from fish too young or too old, or fish that did not grow properly or those that presented the early stages of a disease, led to very low performance in sperm activation or fertilization.

We achieved the best results when using fish 9-11 weeks of age. Even though the natural breeding in this species ensues prior to this age, the gonads largely grow in size between week 6 to week 10, allowing a greater amount of collectable sperm. Also females produce significantly more eggs at week 10 compared to week 6, probably because they are bigger and can store more in their belly.

It is important to emphasize that it was not possible to extract sperm by squeezing the male’s abdomen as for other fish species. Instead, the gonads need to be extracted after male sacrifice and immediately spun vigorously into the extender solution to allow sperm release. This practice led to a lower percentage of mature sperm in the solution since also immature sperm and other kinds of cells were released in the mix. These other not-activatable cells were counted during the analysis since they were not distinguishable from sperm cells, thus underestimating the effective activation yield results. Optimizing the sperm extraction from living males could potentially increase the sperm activation percentages as well as the fertilization success.

In these studies, fertilization were performed with an average of 60.000 sperm per ul. We did not systematically modulate sperm concentration as a variable to maximize the fertilization efficiency. Nevertheless, during our tests, several sperm concentrations arose often due to different gonad dimension or to different volumes of the extender, yet we never observed remarkable differences in sperm activation. We suggest therefore that within a range between 15,000 −200,000 sperm/ul, fertilization occurs at comparable rates.

Another technical point is the use of eppendorf tubes versus cryovials. We found that the conical 1.5ml eppendorf tubes ensured rapid sperm thaws and a better distribution of the eggs and sperm at the bottom, maximizing the fertilization efficiency. Because of their shape, cryovials are less practical since the sperm needs more time to thaw and it is difficult to visualize the exact moment of complete thaw, thus preventing the precise timing needed for subsequent steps of fertilization. Importantly, we revived sperm frozen in Eppendorf tubes from 1 day to a maximum of 4 months, without noticing any remarkable difference in yields of activation (data not shown). Further studies should clarify the long term cryopotential of this setup.

Lastly, even though FBS and BSMIS gave comparable results in terms of sperm activation and fertilization, in our hands FBS resulted in more overall resilience. Notably, FBS gave better results in preserving sperm even when the fish quality was not optimal or when the procedure was not performed with the highest accuracy.

In sum, our optimized protocol and incidental observations should yield fairly consistent results for sperm preservation and IVF, greatly facilitating the husbandry and use of *N. furzeri* as a model organism.

## Materials and Methods

### Fish husbandry and sample collections

All adult fish used were *Nothobranchius furzeri* belonging to the GRZ strain and were raised singularly in 2.8L tanks from the second week of life. Water parameters were pH: 7-8; Kh: 3-5; T: 27 °C. 10% of the water in the system was automatically replaced with clean water every day. Fish were raised in 12 hours of light and 12 hours of darkness. Fish were fed with chironomus twice a day and “Premium Artemia Coppens^®^” twice a day. Natural breeding events occured in 8L tanks with 1-2 males and 3-5 females. To start the breeding event, one box (9cm x 9cm x 4cm) half full of river sand (**ø** < 0,2mm) was put on the bottom of the tank. The boxes were put in the fish tanks for 2-3 hours and the average number of eggs layed by each female in the sand was between 5 and 30. All the fish used for the experiments were from 9 to 11 weeks old.

### Gonads extraction and sperms collection

Males were anesthetized with a tricaine methanesulfonate solution (Sigma, 0.5 mg/ml) for 10+ minutes, until movements and breathing stopped. Fish were dried using tissue paper and decapitated. The belly of the fish was cut open by scissor, the organs removed and the gonads at two sides of the swimming bladder were picked out gently using forceps. Gonads were placed in an Eppendorf tube containing 500 μl of extender solution, grabbed tightly with forceps and and spun thoroughly back and forth for 1 min. After the sperm was released in the solution the gonads were removed from the eppendorf. On average around 30 million sperm (between 7,5 million and 100 million) were released.

### Fresh solutions trials

An individual gonad was placed in 250μl of solution (such as tank water, DE water, BSMIS, HBSS, FBS, Iwamatsu solution) at a certain dilution (1:1, 1:2, 1:4, 1:8) and the sperm was released inside by shaking for 5-10 seconds. 10μl of each solution+sperm mix were placed in a haemocytometer chamber immediately after and the activation of sperm, visualized and recorded. Each trial combination was repeated at least 3 times.

### Extended solution sperm activation trials

One gonad was extracted and shook in extender solutions as previously described. One volume (3μl) of sperm-extender mixture was mixed with two (6μl) or nine volumes (27μl) of activator solutions and immediately transferred to the haemocytometer chamber under the microscope for video recording. Each trial combination was repeated at least 3 times.

### Imaging and video acquisition

For sperm visualization we used an Imager Z1 microscope from Zeiss and for video recording an Axiocam 506 mono, binning 3×3, resolution 2752×2208 and the ZEN 2.3 pro software. Each video was recorded for 8 seconds.

### Imaris conversion and sperm motility analysis

Videos were imported in Imaris 8.1 and frame time points rescaled by 10 times. Black and white colors were inverted in order to have white dots over black background. Particle tracking function was set to identify all sperm with diameter of ±8.6μm. Movements were tracked with autoregressive motion algorithm function, set with max. distance: 25 μm and max. gap size: 5 as parameters. Statistics relative to track displacement length were exported as excel file and analyzed. As a cutoff we set a displacement length of >15μm to distinguish active directionally traveling sperm from not acting or vibrating ones. In case of general drift of the whole sample size we increased the cutoff to >35μm or >50μm (in case of very strong drift).

### Cryoprotectants trials

Two gonads were extracted from a male and were shook into 500μl of a different extender solution (FBS or BSMIS). These mixes were aliquoted in smaller volumes (60μl) and different cryoprotectants (DMSO (Sigma), DMF (Carl Roth), Methanol (Optima), DMA (Sigma) or glycerol) were added in different concentrations in each different eppendorf and immediately mixed. 3μl from each sample were taken and mixed with an activation solution to check the sperm mobility, following the procedure previously described. The rest of each mix was frozen following the cryopreservation procedure. Survivability of the sperm after one-two weeks was checked upon thawing (see below).

### Cryopreservation

Extender-sperm-cryoprotectant mixtures were incubated in 1.5ml eppendorfs for 1 hour at 4°C. The mixtures were distributed in Eppendorf tubes at a volume of 60μl each, the tubes were closed and gently placed at the bottom of a glass beaker sitting in a styrofoam box filled up to 10cm with dry ice. After 15 minutes, eppendorf tubes were quickly moved into a rack in a nitrogen tank, not contacting directly the nitrogen but exposed only to nitrogen vapors for 30 minutes. As the final step the samples were moved directly into liquid nitrogen and stored for days to months.

### Cryopreservation trials

Both gonads from the same individual were placed in 500μl FBS solution and swung for sperm release as previously described. 60μl of the resulting mix were mixed half with 10% DMSO and half with 10% methanol in Eppendorf tubes and incubated for 1 hour at 4°C. The eppendorfs were distributed among Mr. frosty container (−10°C per min), a beaker surrounded with dry ice (−20°C per min), directly on dry ice (−50°C per min), directly in nitrogen gas phase (−100 °C per min), directly in nitrogen liquid (−200°C per min) phase for 30 minutes, until completely frozen. The samples were placed in the nitrogen liquid phase after they reached a temperature of below −50°C. The thawing and monitoring process were performed as previously described.

### Thawing and sperm revival

Frozen samples were thawed directly from liquid nitrogen into a 30°C warm water bath until any ice disappeared. The procedure usually took 1 minute or less. One volume (3μl) of thawed sperm mixtures was mixed with two volumes (6μl) of BSMIS 1:4 on haemocytometer chamber to check for mobility and activation rates.

### *In vitro* fertilization by thawed sperms

Females were quickly anesthetized in a Tricaine methanesulfonate solution (0.5mg/ml) and carefully dried using towel paper to prevent any residual water on the fish surface. Eggs were extracted from the female by gently massaging and slightly pushing their belly. Eggs were laid over a glove and collected through a forceps over the side of a freshly thawed eppendorf with sperm-extender (60μl). A pipette tip was used to push the eggs into the sperm mixture and the eppendorf was gently flicked for 15-20 seconds to prevent egg clumps. 120μl of activation solution was added to the mixtures and mixed flicking the eppendorf for 15-20 seconds. The eggs were incubated in the mixture at room temperature for 10 mins and during the incubation 15ul from the mixture was monitored under the microscope to check the sperm activity. The eggs were finally transferred to a petri dish filled with tank water and the water was replaced twice to wash away any residual cryoprotectant. The petri dish were incubated at 28°C and the eggs were monitored under a microscope for any morphological changes for 4 days.

### Graphs production images acquisitions and enhancement

Raw data depicting track displacement length realized with Imaris were exported as an excel chart. Data relative to all the experiments were pulled in a unique excel file, containing 3 replicates for each individual trial or combination. Total data were used to calculate averages, standard deviations and standard errors, in case of mean of the means. Graphs were produced from these data using excel.

### Images and graphics acquisitions and enhancement

Tracking images were acquired from videos using Imaris snapshot function and brightfield images were acquired using a Leica M80 microscope equipped with a Leica MC170 HD camera. Images were enhanced in brightness, contrast and saturation using GIMP to improve the visual quality.

Graphics and drawings were realized using paint, GIMP and power point.

## Abbreviations used

HBSS: Hank’s Balanced Salt Solution
BSMIS: Buffered sperm motility-inhibiting solution
FBS: Fetal bovine serum
DMSO: Dimethyl sulfoxide
DMF: Dimethylformamide
MetOH: Methanol
DMA: Dimethylacetamide
IVF: *In vitro* fertilization

## Acknowledgements

Imaging analyses were performed in the FACS & Imaging Core Facility at the Max Planck Institute for Biology of Ageing (Cologne).

## Supporting information

**S1 File. IVF protocol.** This is the protocol that we suggest and that uses FBS as extender, DMSO 10% as cryoprotectant and BSMIS 1:4 as activator.

